# Molecular grammar of RNA m^6^A modification in ribonucleoprotein granules

**DOI:** 10.64898/2026.05.26.727807

**Authors:** Mengmeng Xu, Yuchen Li, Tianyu Wang, Jun Liu, Zhi Qi

**Affiliations:** Tsinghua-Peking Center for Life Sciences, Tsinghua University, Beijing 100084, China; Center for Quantitative Biology, Academy for Advanced Interdisciplinary Studies, Peking University, Beijing 100871, China; Peking-Tsinghua Center for Life Sciences, Academy for Advanced Interdisciplinary Studies, Peking University, Beijing 100871, China; State Key Laboratory of Gene Function and Modulation Research, School of Life Sciences, Peking University, Beijing 100871, China; Beijing Advanced Center of RNA Biology (BEACON), Peking University, Beijing 100871, China; School of Physics, Peking University, Beijing 100871, China

## Abstract

Diverse RNA chemical modifications collectively regulate nearly all aspects of RNA metabolism. Among these, *N*^6^-methyladenosine (m^6^A) is the most prevalent internal modification in eukaryotic messenger RNAs. Recent studies have implicated m^6^A in the formation and regulation of ribonucleoprotein (RNP) granules—biomolecular condensates that organize RNA metabolism in space and time. However, the underlying biophysical mechanisms remain to be elucidated. Here, building on SMART, a single-molecule platform for quantitatively measuring RNP granule assembly across nanometer-to-mesoscale regimes, we develop SMART-epi to enable precise control of the m^6^A modification ratio. Using this approach, we show that physiological m^6^A levels inhibit YTHDF2 RNP granule assembly while directing specific RNA-RNA interaction architectures. Moreover, YTHDF1/2/3 exhibit distinct assembly kinetics, suggesting functional divergence. Together, these findings reveal a molecular grammar by which m^6^A modulates condensate dynamics and suggest that RNA modifications can encode regulatory information to tune RNP granule behavior, with potential implications for therapeutic intervention.

## Introduction

RNA plays essential roles in living cells not only through its sequence information but also through extensive chemical modifications ^1^. To date, more than 170 distinct RNA modifications have been identified, collectively regulating nearly all aspects of RNA metabolism, including splicing, localization, stability, translation, and decay ^2^. Among these modifications, *N*^6^-Methyladenosine (m^6^A) is the most prevalent post-transcriptional modification of eukaryotic messenger RNA (mRNA) and long noncoding RNA ^3^. m^6^A functions as a dynamic regulatory mark that recruits specialized RNA-binding proteins (RBPs) to modulate multiple stages of mRNA fate. For example, the YTHDF family of m^6^A reader proteins—comprising the closely related cytoplasmic paralogs YTHDF1, YTHDF2, and YTHDF3—recognizes m^6^A-modified mRNAs through a conserved YTH domain and mediates diverse biological outcomes ^4^. The discovery that m^6^A is reversible and broadly distributed across the transcriptome has driven rapid expansion of the field of epitranscriptomics ^5^.

Most previous studies have focused on detecting RNA modifications and characterizing RNA-RBP interactions at the nanometer scale. However, the underlying biophysical mechanisms by which m^6^A modifications regulate RNA function remain to be elucidated. Transcriptome-wide mapping studies show that, under physiological conditions, the abundance of m^6^A relative to adenosine—defined here as the m^6^A modification ratio ([m^6^A] / [A])—ranges from approximately 1 : 140 to 1 : 500 ^6-9^, indicating that m^6^A is a relatively sparse modification. Biochemical measurements also indicate that m^6^A enhances the RNA-binding affinity of YTHDF family proteins by only about tenfold ^10^. Moreover, the three YTHDF proteins—YTHDF1, YTHDF2, and YTHDF3—exhibit highly similar biochemical properties at the nanometer scale ^11^. These observations raise several important questions. How can such a low- abundance RNA modification, coupled with relatively modest changes in binding affinity, produce substantial regulatory effects in cells? And how do YTHDF family members, despite their similar biochemical properties, execute distinct functional roles in vivo?

Recently, RNA has been demonstrated to be essential for the formation and maintenance of ribonucleoprotein (RNP) granules ^12^. Ribonucleoprotein (RNP) granules ^12^, such as stress granules, processing bodies (P-bodies), and the nucleolus, are biomolecular condensates composed of diverse RNAs and RBPs that play central roles in RNA metabolism. These multicomponent assemblies arise through networks of weak, multivalent interactions, including RNA-RBP binding as well as homotypic and heterotypic interactions among RBPs and RNAs. Dysregulation of RNP granule dynamics has been implicated in numerous human diseases, highlighting their fundamental importance in cellular physiology.

m^6^A modification has been implicated in the formation and regulation of RNP granules. YTHDF1/2/3 can form biomolecular condensates both in vitro and in cells. The resulting mRNA-YTHDF complexes are dynamically partitioned into distinct endogenous RNP granules, such as P-bodies and stress granules, depending on cellular conditions. Upon compartmentalization, the fate of mRNA is regulated in a context-dependent manner by these specific assemblies. For example, under stress conditions, hypermethylated mRNAs are preferentially enriched in stress granules, leading to reduced translation efficiency; this enrichment scales with the number of m^6^A sites. Knockdown of endogenous YTHDF1 or YTHDF3 markedly impairs stress granule formation and abolishes the enrichment of both methylated and unmethylated mRNAs within these compartments ^13-15^.

RNP granules represent complex biological systems characterized by cross-scale emergence, in which biological functions arise from molecular interactions spanning multiple length scales. These scales range from nanometer-scale biomolecular interactions to mesoscale organization, where RNP granules form spatially and temporally defined intracellular structures that execute emergent biological functions. Importantly, the assembly and disassembly of RNP granules are highly dynamic and strongly pathway dependent, such that the composition of a granule at any given time is shaped by its recent assembly history and, in turn, constrains its subsequent compositional evolution.

To understand how the biological functions of RNP granules emerge from higher-order molecular assemblies, our previous work ^16^ established an integrated experimental and theoretical framework to quantitatively investigate RNP granule dynamics, with a particular focus on assemblies formed on long RNA substrates. By leveraging the precise controllability of in vitro single-molecule approaches based on the high-throughput DNA Curtains platform ^17-19^, we developed a single-molecule method termed SMART (group <u>s</u>ingle-<u>m</u>olecule <u>a</u>ssay for <u>r</u>ibonucleoprotein granules). This approach enables direct and quantitative measurements of RNP granule assembly and disassembly kinetics across molecular-to-mesoscale regimes.

In this work, to further investigate the molecular mechanisms by which m^6^A regulates RNP granule assembly, we developed an upgraded version of this method, SMART-epi (group <u>s</u>ingle-<u>m</u>olecule <u>a</u>ssay for ribonucleoprotein granules in <u>epi</u>transcriptomics), which allows precise control of the m^6^A modification ratio ([m^6^A] / [A]). Using SMART-epi, we examined RNP granules composed of either a single YTHDF family member (single-component granules) or two members (two-component granules). Our results demonstrate that physiological m^6^A ratio ([m^6^A] / [A] = 1 : 400), which is low-abundance, inhibits the assembly kinetics of YTHDF2 RNP granules while simultaneously guiding the formation of specific RNA-RNA interaction architectures within these condensates. We further show that YTHDF1, YTHDF2, and YTHDF3 exhibit distinct assembly kinetics. In mixed systems, YTHDF1 acts as a kinetic inhibitor that attenuates YTHDF2 assembly, whereas YTHDF3 functions as a kinetic activator that promotes YTHDF2 assembly.

Together, our results show that, in addition to RNA-protein, protein-protein, and RNA-RNA interactions, low-abundance of m^6^A modification ratio also modulates the cross-scale kinetics of RNP granule assembly. These findings suggest that m^6^A-dependent regulation may provide a means to precisely tune RNP granule dynamics, potentially enabling targeted manipulation of condensate behavior and opening new avenues for therapeutic intervention in human disease.

## Results

### Developing SMART-epi to study the molecular mechanism of RNA m^6^A modification

We first inserted a single T7 promoter into Lambda DNA and introduced these DNA substrates into a microfluidic flow chamber of the DNA Curtains platform ^17-19^. Individual DNA molecules were tethered to supported lipid bilayers through specific streptavidin–biotin interactions (Fig. 1a(i) and Supplementary Fig. 1). T7 RNA polymerase (T7 RNAP) together with a mixture of nucleoside triphosphates (NTPs), including Fluor647X-labeled UTP and *N*^6^-methyl-ATP, was then introduced to initiate in vitro transcription. The resulting nascent RNA transcripts, generated from identical DNA templates, were visualized as localized Fluor647X-enriched regions downstream of the T7 promoter (Fig. 1a(ii) and b(i)). Incorporation of *N*^6^-methyl-ATP produced RNA transcripts containing m^6^A modifications. By adjusting the concentration of *N*^6^-methyl-ATP, we controlled the m^6^A modification ratio ([m^6^A] / [A]) under defined experimental conditions. The resulting m^6^A modification ratios were calibrated by mass spectrometry (Supplementary Fig. 2).

**Figure 1.**
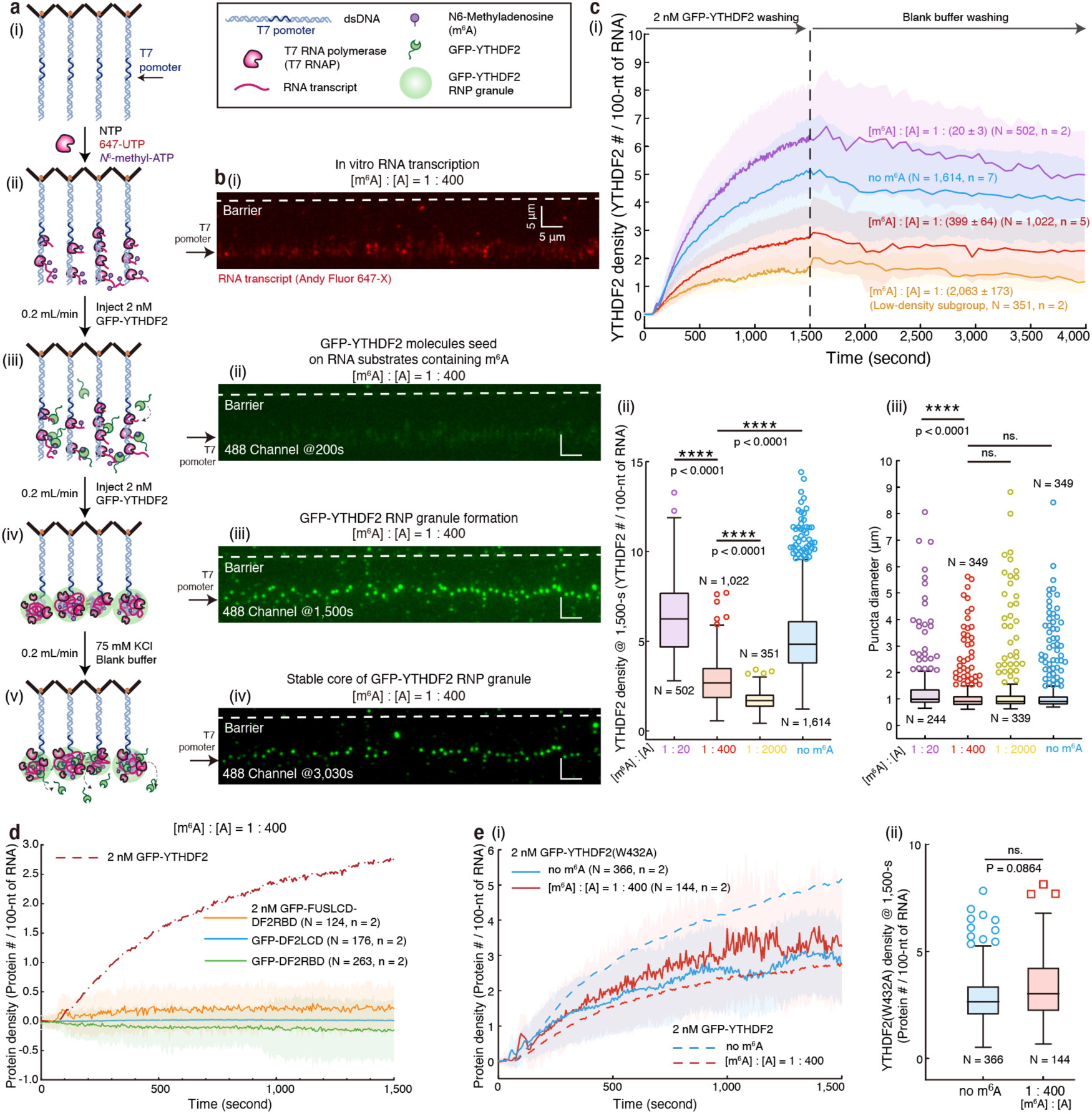
SMART-epi enables quantitative analysis of m^6^A-dependent RNP granule dynamics. (**a**) Schematic of the SMART-epi platform for quantifying YTHDF2 RNP granule assembly and disassembly kinetics across defined m^6^A modification ratios ([m^6^A] / [A]). (**b**) Representative wide-field total internal reflection fluorescence microscopy (TIRFM) images acquired during SMART-epi measurements with 2 nM GFP-YTHDF2. Panels (ii)-(iii) show the assembly phase, followed by buffer exchange to initiate disassembly (iii)-(iv). Panel (i) shows corresponding RNA fluorescence. (**c**) (i) Assembly kinetics of YTHDF2 RNP granules under varying m^6^A ratios ([m^6^A] / [A] = 0, 1 : 2,000, 1 : 400, 1 : 20). Data represent mean ± s.d.; (ii)-(iii) Box plots of assembly density (ii) and puncta diameter (iii) at 1,500-s. (**d**) Assembly kinetics of YTHDF2 truncation variants (GFP-FUSLCD-DF2RBD, GFP-DF2RBD, GFP-DF2LCD) compared with full-length GFP-YTHDF2 ([m^6^A] / [A] = 1 : 400). (**e**) (i) Assembly kinetics of GFP-YTHDF2(W432A) under m^6^A-deficient and m^6^A-containing conditions, with wild-type controls. (ii) Box plot of assembly density at 1,500-s. N indicates total puncta pooled across independent experiments (n). The function of “boxplot” in GraphPad Prism software (Version 8.0.2) was used to plot the boxplots. For each boxplot, the center line within the box represented the median. The bottom edge of the box represents 25^th^ percentile, and the top is 75^th^ percentile. Most extreme data points are covered by the whiskers except outliers. The “ο” symbol is used to represent the outliers. Unpaired two-tailed t-tests were used to analyze the significant differences between the two groups of data. Unpaired (one-way) ANOVA was used to examine the statistical significance across the multiple groups. P-value style was as follows: GP: 0.1234 (ns), 0.0332 (*), 0.0021 (**), 0.0002 (***), <0.0001 (****). The confidence level was set at 95% for all analyses. Data are presented as mean ± s.d. The working buffer for SMART is 40 mM Tris-HCl (pH 7.5), 75 mM KCl, 0.5 mg/mL BSA, 2 mM MgCl_2_, and 1 mM DTT, and 8 unit/mL RNasin® Plus RNase Inhibitor.

We first examined a physiological condition ([m^6^A] / [A] = 1 : 400). After m^6^A-containing RNA substrates were generated within the flow chamber, 2 nM GFP-tagged YTHDF2 (GFP-YTHDF2; Supplementary Fig. 3) was introduced into the flow cell for 1,500 seconds. We focused initially on YTHDF2 because it is relatively abundant across many tissues ^3^. Following protein introduction, we observed the progressive growth of green fluorescent puncta downstream of the T7 promoter (Fig. 1a(iii)-(iv), b(ii)-(iii)), indicating the formation of YTHDF2 RNP granules ^16^.

The chamber was subsequently washed with protein-free working buffer for an additional 3,600 seconds (Fig. 1a(iv)-(v), b(iii)-(iv)). Quantitative analysis of total nucleotides (nts) of RNA transcripts in each granules, inferred from Fluor647X fluorescence intensity, yielded an average transcript length of (3.9 ± 3.3) × 10^5^ nts (mean ± s.d., N = 1,022). Mass spectrometry measurements confirmed an m^6^A modification ratio of 1 : (399 ± 64) (mean ± s.d., n = 6; Supplementary Fig. 2(iv)), corresponding to an estimated total of 244 ± 207 m^6^A sites per transcript.

Using SMART ^16^, we quantified protein density—defined as the number of bound proteins per 100-nt of RNA—as a function of time (red curve in Fig. 1c(i)). At 1,500 seconds, YTHDF2 RNP granules reached a protein density of 2.7 ± 1.1 (mean ± s.d., N = 1,022; [m^6^A] / [A] = 1 : 400; Fig. 1c(ii) and Supplementary Fig. 4c), corresponding to an average of (1.06 ± 0.97) × 10^4^ YTHDF2 molecules per granule. The mean granule diameter was 1.1 ± 0.8 μm (mean ± s.d., N = 349; [m^6^A] / [A] = 1 : 400; Fig. 1c(iii)). After a 1-hour wash with protein-free buffer, only ∼19% of YTHDF2 molecules dissociated from the granules. Accordingly, the protein density at 5,100 seconds remained high (2.2 ± 1.1; mean ± s.d., N = 1,022; [m^6^A] / [A] = 1 : 400), indicating that YTHDF2 RNP granules form a stable core architecture. We refer to this upgraded approach as SMART-epi, which enables direct measurement of RNP granule assembly on long RNA substrates containing controlled m^6^A modifications.

### A specific DF2 LCD governs the assembly kinetics of YTHDF2 RNP granules

To determine how individual domains of YTHDF2 contribute to granule formation, we next dissected the domain architecture of YTHDF2. YTHDF2 contains two principal regions (Supplementary Fig. 3a(i)): a low-complexity domain (DF2LCD) and an RNA-binding domain (DF2RBD).

We purified DF2RBD (Supplementary Fig. 5a-b) and DF2LCD (Supplementary Fig. 6a-b) separately. Electrophoretic mobility shift assays (EMSAs) showed that DF2RBD or DF2LCD binds a 34-nt RNA substrate containing a single m^6^A modification with affinities of 602 ± 194 nM (DF2RBD, mean ± s.d., n = 3, Supplementary Fig. 5c), or 719 ± 103 nM (DF2LCD, mean ± s.d., n = 3; Supplementary Fig. 6c), respectively—substantially weaker than wild-type YTHDF2 (79 ± 14 nM, mean ± s.d., n = 3, Supplementary Fig. 3c). These results indicate that each domain binds RNA only weakly in isolation, whereas cooperative interactions between the LCD and RBD enhance RNA recognition in YTHDF2.

SMART-epi measurements further revealed that neither GFP-DF2RBD nor GFP-DF2LCD assembles into RNP granules on RNA substrates containing a physiological m^6^A ratio (green and blue curves in Fig. 1d, and Supplementary Fig. 5e and 6e). Compared with the assembly kinetics of YTHDF2 (red dashed line in Fig. 1d), these observations suggested that the LCD may play a critical role in governing the assembly kinetics of YTHDF2 RNP granules.

To test this hypothesis, we generated a chimeric protein in which the DF2LCD was replaced by the well-characterized LCD of human FUS (FUSLCD-DF2RBD) (Supplementary Fig. 7a-b). The FUS LCD has been extensively studied for its condensation properties ^20,21^. Unlike DF2RBD alone, 20 μM GFP-FUSLCD-DF2RBD formed condensates in vitro, and the addition of 34-nt RNA substrates containing either zero or one m^6^A modification further increased condensate size (Supplementary Fig. 7d). EMSAs showed that the chimeric protein exhibited intermediate RNA-binding affinity (437 ± 257 nM, mean ± s.d., n = 3; Supplementary Fig. 7c), stronger than DF2RBD but weaker than YTHDF2. Strikingly, however, SMART-epi measurements revealed that FUSLCD–DF2RBD failed to assemble into RNP granules on RNA substrates containing a physiological m^6^A ratio (orange curve in Fig. 1d). Together, these results demonstrate that a specific LCD is required to control the assembly kinetics of YTHDF2 RNP granules.

### The m^6^A modification ratio regulates the assembly and disassembly kinetics of YTHDF2 RNP granules

Having established the assembly and disassembly kinetics of YTHDF2 RNP granules under physiological conditions, we next examined whether the m^6^A modification ratio modulates these dynamics. We performed SMART-epi measurements using RNA substrates lacking m^6^A modifications (Supplementary Fig. 4a(i)), with the resulting kinetics shown as the blue curve in Fig. 1c(i). Surprisingly, in the absence of m^6^A, the YTHDF2 protein density at 1,500 s increased dramatically from ∼2.7 under physiological conditions to 5.1 ± 1.9 (mean ± s.d., N = 1,614; [m^6^A] / [A] = 0; Fig. 1c(ii) and Supplementary Fig. 4c). By contrast, the granule diameter (1.3 ± 1.0 μm; mean ± s.d., N = 349; [m^6^A] / [A] = 0) remained comparable to that observed under physiological conditions (Fig. 1c(iii)), whereas the total number of YTHDF2 molecules per granule ((1.99 ± 1.16) × 10^4^, mean ± s.d., N = 1,614, [m^6^A] / [A] = 0) increased by approximately twofold.

Under physiological conditions ([m^6^A] / [A] = 1 : 400), the RNA substrates are approximately 390,000 nts in length and contain only ∼240 m^6^A-modified adenosines. Despite their low abundance, these m^6^A modifications exert a strong inhibitory effect on YTHDF2 granule assembly, revealing an unexpected regulatory role for sparse RNA modifications.

To validate the SMART-epi measurements, we purified GFP-YTHDF2(W432A) (Supplementary Fig. 8), a mutant with markedly reduced m^6^A recognition capacity ^10^. SMART-epi analysis showed that YTHDF2(W432A) exhibited nearly identical RNP granule assembly kinetics under both the m^6^A-free condition and the physiological condition (Fig. 1e). These results demonstrate that specific recognition of m^6^A directly modulates the assembly kinetics of YTHDF2 RNP granules and further confirm the specificity and robustness of the SMART-epi assay.

We next asked which m^6^A modification ratio produces the strongest inhibitory effect on YTHDF2 RNP granule assembly. To address this question, we reduced the m^6^A modification ratio to 1 : (2,063 ± 173) (mean ± s.d., n = 6; Supplementary Fig. 2(iv)). Under this condition, SMART-epi measurements revealed that YTHDF2 RNP granule assembly segregated into two distinct kinetic subpopulations: a low-density subgroup and a high-density subgroup (Supplementary Fig. 4b(i)-(ii)). At 1,500 seconds, the high-density subgroup reached a YTHDF2 density of 3.4 ± 0.8 (mean ± s.d., N = 952), slightly exceeding the density observed under physiological conditions ([m^6^A] / [A] = 1 : 400). In contrast, the low-density subgroup displayed a markedly reduced YTHDF2 density of 1.7 ± 0.5 (mean ± s.d., N = 351), substantially lower than that observed at the physiological m^6^A ratio (Supplementary Fig. 4b(iii)-(v) and c).

These observations indicate the existence of an m^6^A density threshold that governs YTHDF2 RNP granule assembly. When the m^6^A ratio falls below this threshold, a subset of assemblies recovers high-density behavior comparable to that observed in the absence of m^6^A, whereas another subset remains strongly inhibited. For subsequent analysis, we used the low-density subgroup to represent the effective assembly behavior at [m^6^A] / [A] = 1 : 2,000 (the orange curve in Fig. 1c(i)). Under this condition, the RNA substrates are approximately 760,000 nucleotides in length and contain only ∼92 m^6^A-modified adenosines. Despite their extremely low abundance, these sparse m^6^A modifications exert a disproportionately strong inhibitory effect on YTHDF2 RNP granule assembly.

Finally, we examined the effects of an artificially elevated m^6^A modification density. Increasing the m^6^A modification ratio to 1 : (20 ± 3) (mean ± s.d., n = 6; Supplementary Fig. 2(iv)), well above physiological levels, markedly enhanced YTHDF2 assembly on RNA substrates (purple curve in Fig. 1c(i)). Accordingly, the YTHDF2 protein density at 1,500 s increased from ∼5.1 under the unmethylated condition to 6.3 ± 1.9 (mean ± s.d., N = 502; [m^6^A] / [A] = 1 : 20; Fig. 1c(ii) and Supplementary Fig. 4a(ii) and 4c). These results indicate that supraphysiological m^6^A densities strongly promote YTHDF2 RNP granule assembly.

Our previous study ^16^ using SMART revealed that RNP granules can establish distinct pathway-dependent RNA-RNA interaction architectures. Consistent with this idea, Parker and colleagues recently demonstrated that G3BP1 promotes the formation of intermolecular RNA-RNA interaction networks within biomolecular condensates ^22,23^. To investigate the molecular mechanism by which the m^6^A modification ratio regulates RNP granule assembly, we first examined whether YTHDF2 similarly promotes intermolecular RNA-RNA interaction networks within YTHDF2 RNP granules (Supplementary Fig. 9a).

Following the experimental strategy described previously ^22^, we mixed 10 μM YTHDF2 with 0.5 μM RNA substrates (196-nt) either lacking m^6^A or containing a physiological modification ratio, resulting in the formation of YTHDF2-RNA co-condensates (Supplementary Fig. 9b(i), c(i)). After the addition of proteinase K, the fluorescence signal of YTHDF2 rapidly disappeared, confirming efficient protein digestion. In contrast, the fluorescence signal from the RNA substrates decayed much more slowly under both conditions (no m^6^A or [m^6^A] / [A] = 1 : 400) (Supplementary Fig. 9b(ii)-(iii), c(ii)-(iii)).

Interestingly, when wt YTHDF2 was replaced with the m^6^A-recognition mutant YTHDF2(W432A) (Supplementary Fig. 8), both the protein and RNA fluorescence signals disappeared nearly simultaneously following the same proteinase K treatment (Supplementary Fig. 9d-e). These observations suggest that YTHDF2 promotes the formation of intermolecular RNA-RNA interaction networks within YTHDF2 RNP granules, and that the conserved residue W432, which mediates both m^6^A recognition and RNA-binding capacity, plays a critical role in regulating the formation of these RNA-RNA interaction architectures.

We next performed RNase cleavage experiments to further probe the RNA structural organization within the granules. Two enzymes were used: RNase A, which digests both single-stranded RNA (ssRNA) and double-stranded RNA (dsRNA), and RNase I_f_, which specifically cleaves ssRNA. In the absence of YTHDF2, ∼92% of the RNA substrates (no m^6^A or [m^6^A] / [A] = 1 : 400) were cleaved by RNase I_f_ within 780 seconds (1 U/mL RNase I_f_, Supplementary Fig. 10b). In contrast, within YTHDF2 RNP granules, even with a tenfold higher concentration of RNase I_f_ and after 1,740 s, only ∼62% of RNA substrates were cleaved (10 U/mL RNase I_f_, Supplementary Fig. 11b-e). As a control, RNase A treatment resulted in rapid digestion of ∼94% of RNA substrates lacking m^6^A within 15 seconds, both in the absence of YTHDF2 (1 μg/mL RNase A, Supplementary Fig. 10a) and within YTHDF2 RNP granules (1 μg/mL RNase A, Supplementary Fig. 11a and d). These results suggest that assembly of YTHDF2 RNP granules promotes the formation of RNA-RNA interaction architectures enriched in dsRNA structures. Such structures slow the digestion kinetics of the ssRNA-specific enzyme RNase I_f_ but not the broadly active RNase A.

Based on these observations, we propose a molecular mechanism by which the m^6^A modification ratio regulates YTHDF2 RNP granule kinetics. Specifically, the m^6^A modification ratio exhibits a threshold near [m^6^A] / [A] = 1 : 2,000, at which RNA-RNA interaction architectures within YTHDF2 RNP granules reach maximal dsRNA abundance and minimal ssRNA accessibility. Because EMSAs show that YTHDF2 binds ssRNA with higher affinity than dsRNA (Supplementary Fig. 12), this structural reorganization results in maximal inhibition of YTHDF2 RNP granule assembly (Fig. 1c(ii)). Interestingly, even at the physiological m^6^A modification ratio ([m^6^A] / [A] = 1 : 400), RNA substrates exert a measurable inhibitory effect on YTHDF2 assembly compared with unmethylated RNA (Fig. 1c(i)). Together, these findings demonstrate that, in addition to RNA-protein, protein-protein, and RNA-RNA interactions, the m^6^A modification ratio can also modulate the assembly kinetics of YTHDF2 RNP granule. This is the first molecular mechanism of the m^6^A modification ratio.

### Physiological m^6^A levels promote the establishment of specific RNA-RNA interaction architectures within YTHDF2 RNP granules

The previous study ^16^ using SMART demonstrated that protein concentration governs the assembly kinetics of RNP granules. Under m^6^A-free conditions, we first introduced 1 nM YTHDF2 for 1,200 seconds. The assembly kinetics at 1 nM were markedly slower than at 2 nM (Fig. 2a(i)), confirming that protein concentration modulates granule assembly. When the system was subsequently switched to 2 nM YTHDF2 for an additional 1,200 seconds (Fig. 2a(i)), the final YTHDF2 density at 2,400-s was 2.5 ± 0.8 (mean ± s.d., N = 358; Fig. 2a(ii)), substantially lower than that observed under continuous 2 nM exposure for 2,400 seconds (4.3 ± 0.9, mean ± s.d., N = 630; Fig. 2a(ii)). These findings indicate that the RNA-RNA interaction architecture formed under constant 2 nM conditions differs from that generated through stepwise exposure (1 nM followed by 2 nM), revealing pronounced pathway dependence, consistent with the previous reference ^16^.

**Figure 2.**
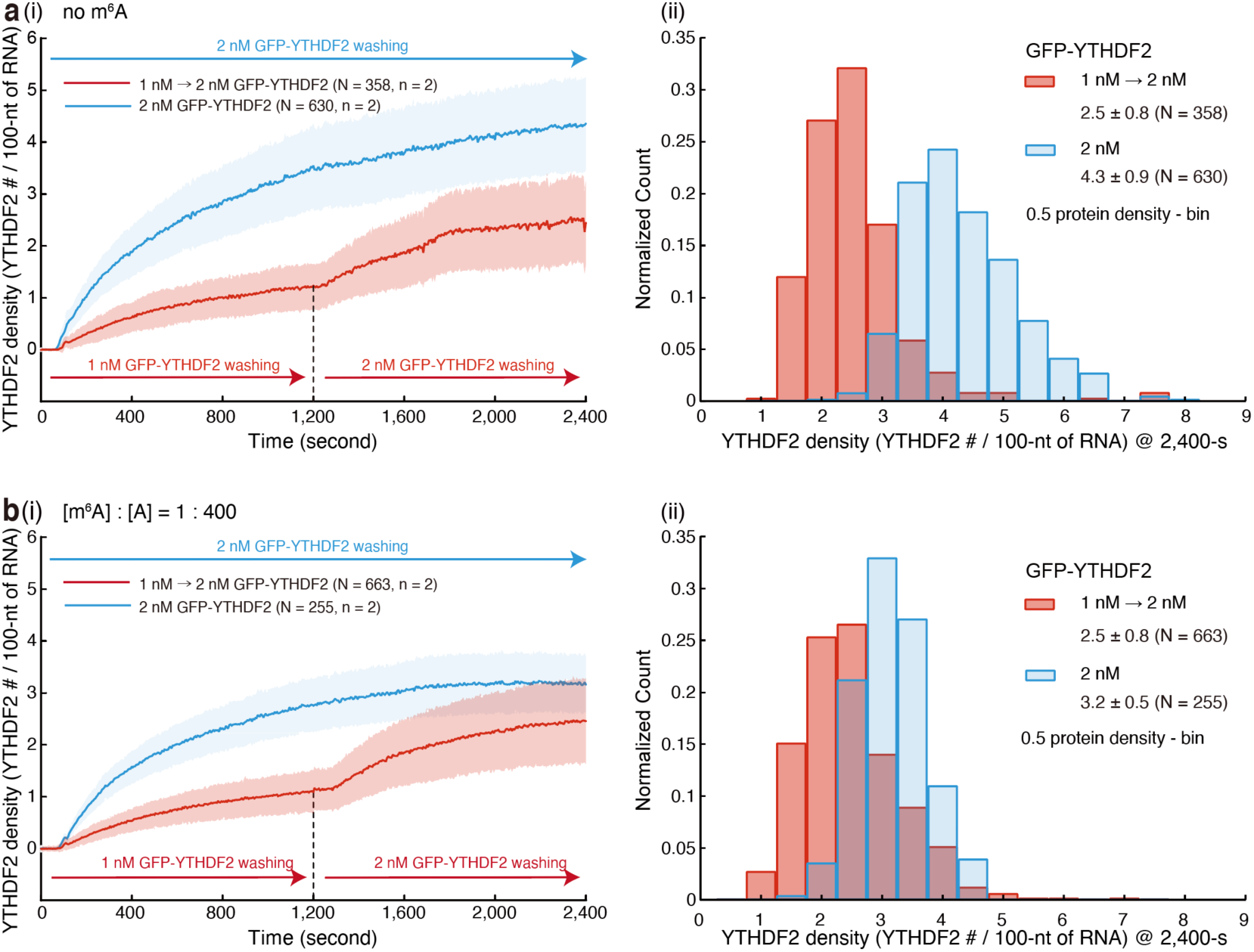
Assembly kinetics under sequential protein exposure. (**a-b**) (i) Assembly kinetics of YTHDF2 measured by SMART-epi under sequential versus constant protein exposure. Red trace: the chamber was first equilibrated with 1 nM YTHDF2 for 1,200 seconds, followed by exposure to 2 nM YTHDF2 for an additional 1,200 seconds. Blue trace: the chamber was continuously exposed to 2 nM YTHDF2 for 2,400 seconds. (ii) Histograms of YTHDF2 assembly density at 2,400 seconds for the conditions shown in (i), with a bin width of 0.5. [m^6^A] / [A] = 0 in (a) and 1 : 400 in (b). Data are presented as mean ± s.d. N denotes the total number of puncta pooled across independent experiments (n).

We next repeated these experiments under a physiological m^6^A modification ratio ([m^6^A] / [A] = 1 : 400). As expected, assembly during the initial 1,200-s remained slower at 1 nM than at 2 nM (Fig. 2b(i)). Surprisingly, however, after switching from 1 nM to 2 nM YTHDF2 for the subsequent 1,200 seconds, the final density at 2,400-s was 2.5 ± 0.8 (mean ± s.d., N = 663; Fig. 2b(ii)), closely approximating that obtained under continuous 2 nM exposure for 2,400 seconds (3.2 ± 0.5, mean ± s.d., N = 255; Fig. 2b(ii)). Thus, under physiological m^6^A conditions, the RNA-RNA interaction architectures generated by distinct assembly pathways converge to a similar final state. On this basis, we propose a second molecular mechanism by which the m^6^A modification ratio regulates YTHDF2 RNP granule kinetics: physiological m^6^A levels act as a guiding framework that promotes the establishment of specific RNA-RNA interaction architectures within YTHDF2 RNP granules for specific biological functions, thereby reducing pathway dependence and constraining the accessible structural states.

### YTHDF family members display distinct assembly kinetics

In addition to YTHDF2, the YTHDF family includes YTHDF1 (Supplementary Fig. 13a(i)) and YTHDF3 (Supplementary Fig. 14a(i)), which share highly similar amino acid sequences, net charge distributions, and hydrophobicity profiles along their lengths ^11^. Following purification of YTHDF1 (Supplementary Fig. 13a(ii)-b) and YTHDF3 (Supplementary Fig. 14a(ii)-b), EMSAs demonstrated that all three YTHDF proteins exhibit comparable RNA-binding affinities toward 34-nt RNA substrates containing a single m^6^A modification (Supplementary Fig. 13c, Supplementary Fig. 14c, and Supplementary Fig. 15a). In vitro droplet assays further showed that each YTHDF protein independently forms condensates, and that addition of m^6^A-modified RNA promotes the formation of larger co-condensates (Supplementary Fig. 15b-d).

Despite these striking biochemical and biophysical similarities, SMART-epi measurements revealed that YTHDF1, YTHDF2, and YTHDF3 exhibit distinct assembly kinetics within RNP granules under both physiological m^6^A conditions ([m^6^A] / [A] = 1 : 400) and m^6^A-free conditions (Fig. 3a(i)-(ii), Supplementary Fig. 13d, and Supplementary Fig. 14d). Specifically, YTHDF2 achieves the highest protein density upon assembly, followed by YTHDF3, whereas YTHDF1 exhibits the lowest protein density. These findings demonstrate that nanoscale similarities in sequence composition and RNA-binding affinity do not necessarily translate into equivalent mesoscale assembly behaviors. Instead, distinct kinetic features emerge from higher-order molecular assembly, reflecting a cross-scale emergence of function. Such kinetic divergence may enable temporally differentiated biological functions of YTHDF proteins during RNP granule formation and remodeling in living cells.

**Figure 3.**
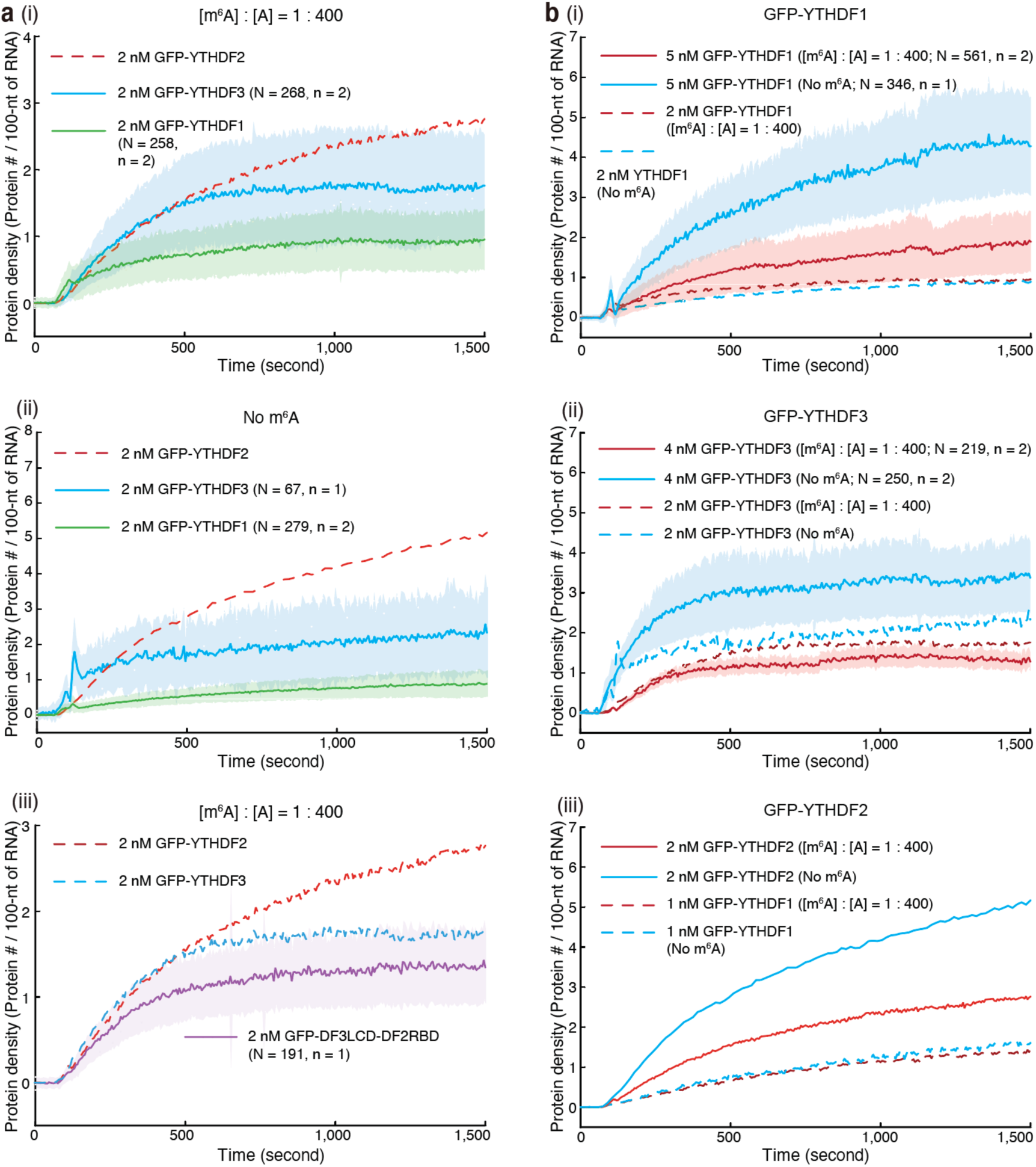
YTHDF family members display distinct assembly kinetics. (**a**) (i)-(ii) Assembly kinetics of YTHDF family members measured by SMART-epi. Green trace, YTHDF1; blue trace, YTHDF3; red dashed trace, YTHDF2. [m^6^A] / [A] = 1 : 400 in (i) and 0 in (ii). (iii) Assembly kinetics of the chimeric construct GFP-DF3LCD-DF2RBD compared with full-length GFP-YTHDF2 and GFP-YTHDF3 ([m^6^A] / [A] = 1 : 400). (**b**) Assembly kinetics under varying protein concentrations and m^6^A modification levels. (i) YTHDF1: solid lines, 5 nM protein; dashed lines, 2 nM protein. (ii) YTHDF3: solid lines, 4 nM protein; dashed lines, 2 nM protein. (iii) YTHDF2: solid lines, 2 nM protein; dashed lines, 1 nM protein. Blue, [m^6^A] / [A] = 0; red, [m^6^A] / [A] = 1 : 400. Data are presented as mean ± s.d. N denotes the total number of puncta pooled across independent experiments (n).

The preceding experiments demonstrated that a specific DF2 LCD governs the assembly kinetics of YTHDF2 RNP granules. We therefore asked whether specific LCD identity also contributes to the cross-scale emergence observed among YTHDF family RNP granules. To address this, we generated another chimeric fusion protein in which the LCD of YTHDF3 replaced the LCD of YTHDF2 (DF3LCD-DF2RBD, Supplementary Fig. 16a-b). EMSAs revealed an intermediate RNA-binding affinity (92 ± 110 nM, mean ± s.d., n = 3; Supplementary Fig. 16c), weaker than both wt YTHDF2 (Supplementary Fig. 3c) and YTHDF3 (Supplementary Fig. 13c). Interestingly, SMART-epi measurements (Supplementary Fig. 16d) showed that the assembly kinetics of DF3LCD-DF2RBD RNP granules closely resemble those of YTHDF3 rather than YTHDF2 (Fig. 3a(iii)). These findings demonstrate that specific LCD identity is a key determinant of mesoscale assembly kinetics and underlies the cross-scale emergence observed within the YTHDF protein family.

In Fig. 3a(i)-(ii), SMART measurements revealed that at 2 nM, YTHDF1 and YTHDF3 exhibit nearly identical RNP granule assembly kinetics under both m^6^A-free conditions and physiological m^6^A conditions (Fig. 3b(i)-(ii)), similar to the behavior observed for 1 nM YTHDF2 (Fig. 3b(iii) and Supplementary Fig. 17). Interestingly, upon increasing protein concentration—YTHDF1 from 2 nM to 5 nM (Fig. 3b(i)), YTHDF3 from 2 nM to 4 nM (Fig. 3b(ii)), or YTHDF2 from 1 nM to 2 nM (Fig. 3b(iii))—the assembly kinetics under m^6^A-free and physiological m^6^A conditions became clearly distinguishable. These findings indicate that YTHDF family members possess distinct concentration thresholds that modulate RNP granule assembly in the presence of physiological m^6^A ratio. Such threshold-dependent behavior further underscores the differential kinetic regulation among YTHDF proteins despite their biochemical similarities.

### Assembly of two-component YTHDF family RNP granules

Endogenous YTHDF-containing RNP granules likely comprise multiple YTHDF family members together. For example, YTHDF3 has been reported to cooperate with YTHDF1 to promote translation of m^6^A-modified mRNAs, and to synergize with YTHDF2 in facilitating mRNA decay ^24^. We therefore investigated how mixtures of two different YTHDF family members assemble on shared RNA substrates containing physiological m^6^A levels ([m^6^A] / [A] = 1 : 400).

We first co-introduced 1 nM GFP-YTHDF2 and 1 nM mCherry-YTHDF1 into the flow cell (Fig. 4a and Supplementary Fig. 18). Under these conditions, YTHDF1 exhibited minimal assembly on RNA substrates over 1,500 seconds (purple solid line, Fig. 4a). Notably, YTHDF2 assembly was also markedly suppressed relative to its single-component behavior (green solid line versus blue dashed line, Fig. 4a). Consequently, the total assembly kinetics of the mixture (yellow solid line, Fig. 4a) were substantially slower than those observed for 2 nM YTHDF2 alone (red dashed line, Fig. 4a). These results indicate that, in the context of mixed YTHDF2-YTHDF1 RNP granules, YTHDF1 acts as a kinetic inhibitor, attenuating YTHDF2 assembly within the RNP granules.

**Figure 4.**
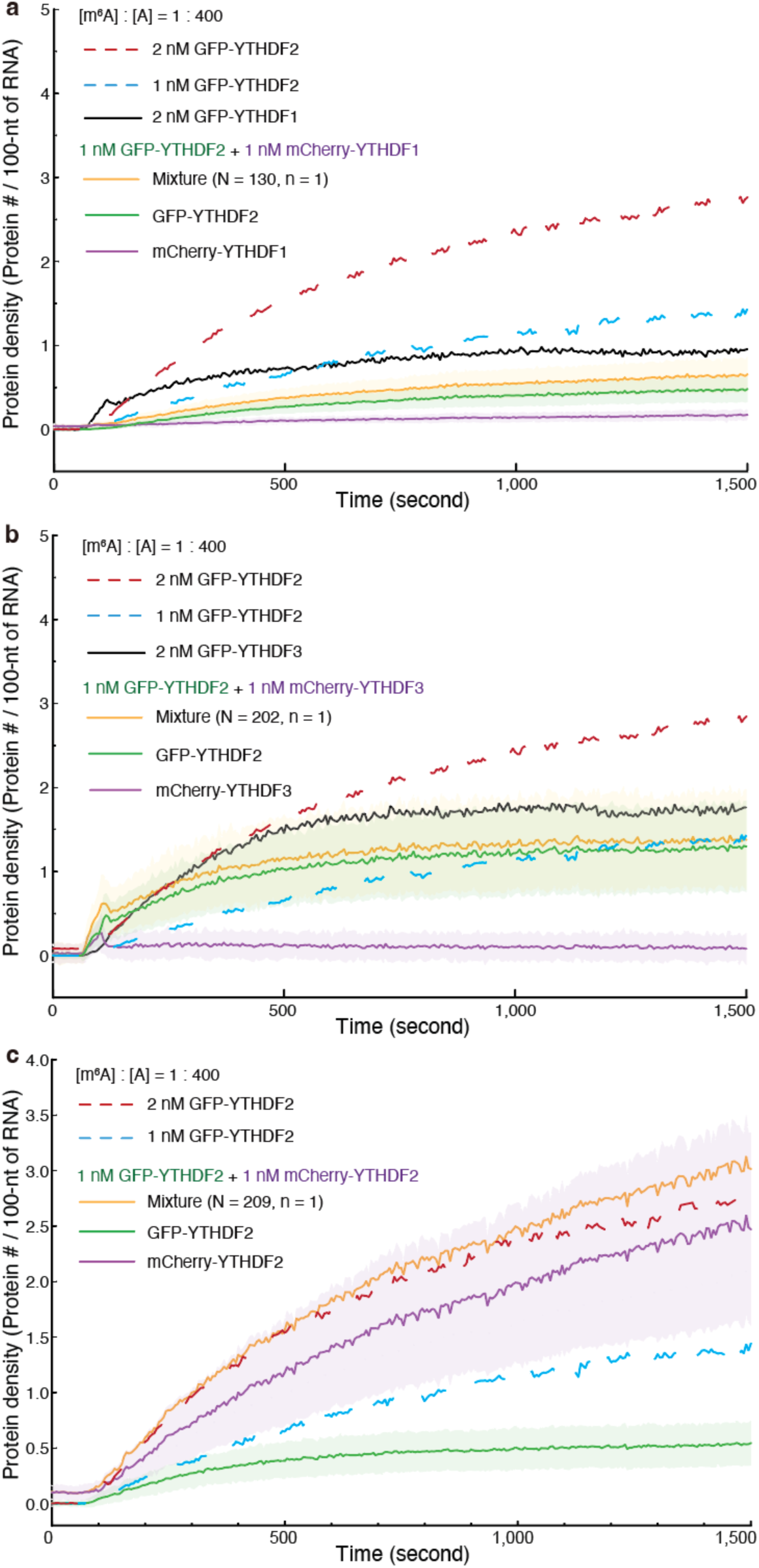
Assembly of two-component YTHDF family RNP granules. (**a**) Assembly kinetics of 1 nM GFP-YTHDF2 and 1 nM mCherry-YTHDF1. Solid orange line, mixture of two proteins; solid green line, GFP-YTHDF2; solid magenta line, mCherry-YTHDF1; solid black line, 2 nM mCherry-YTHDF1. (**b**) Assembly kinetics of 1 nM GFP-YTHDF2 and 1 nM mCherry-YTHDF3. Solid orange line, mixture of two proteins; solid green line, GFP-YTHDF2; solid magenta line, mCherry-YTHDF3; solid black line, 2 nM mCherry-YTHDF3. (**c**) Assembly kinetics of 1 nM GFP-YTHDF2 and 1 nM mCherry-YTHDF2. Solid orange line, mixture of two proteins; solid green line, GFP-YTHDF2; solid magenta line, mCherry-YTHDF2. Dashed red line, 2 nM GFP-YTHDF2; dashed blue line, 1 nM GFP-YTHDF2. [m^6^A] / [A] = 1 : 400. Data are presented as mean ± s.d. N denotes the total number of puncta pooled across independent experiments (n).

We next co-introduced 1 nM GFP-YTHDF2 and 1 nM mCherry-YTHDF3 into the flow cell (Fig. 4b and Supplementary Fig. 19). Under these conditions, similar to YTHDF1, YTHDF3 exhibited minimal assembly on RNA substrates over 1,500 seconds (purple solid line, Fig. 4b). Notably, however, YTHDF2 assembly was markedly enhanced relative to its single-component behavior at 1 nM (green solid line versus blue dashed line, Fig. 4b). Nevertheless, the total assembly kinetics of the mixture (yellow solid line, Fig. 4b) remained slower than those observed for 2 nM YTHDF2 alone (red dashed line, Fig. 4b). These findings indicate that, within mixed YTHDF2-YTHDF3 RNP granules, YTHDF3 functions as a kinetic activator, promoting YTHDF2 assembly, albeit not to the level achieved by doubling YTHDF2 concentration.

As a control, we co-introduced 1 nM GFP-YTHDF2 and 1 nM mCherry-YTHDF2 into the flow cell (Fig. 4c). The total assembly kinetics of the mixed population (yellow solid line, Fig. 4c) closely matched those observed for 2 nM YTHDF2 alone (red dashed line, Fig. 4c).

## Discussion

In this study, building on SMART ^16^, we developed SMART-epi to investigate what are the molecular mechanisms of RNA m^6^A modification in RNP granules. We identified protein concentration (Fig. 3b) and the m^6^A modification ratio (Fig. 1c(i)) as two key parameters that regulate YTHDF-family RNP granule formation. Beyond a critical protein concentration threshold, the physiological m^6^A ratio ([m^6^A] / [A] = 1 : 400) inhibits the assembly kinetics of YTHDF-family RNP granules. We further found that the physiological m^6^A ratio acts as a guiding framework that promotes the establishment of specific RNA-RNA interaction architectures within YTHDF2 RNP granules (Fig. 2), thereby reducing pathway dependence and constraining the range of accessible structural states. In addition, YTHDF1, YTHDF2, and YTHDF3 exhibit distinct assembly kinetics under physiological m^6^A conditions (Fig. 3a(i)-(ii)). These kinetic differences arise from higher-order molecular assembly and reflect the cross-scale emergence of functional behaviors. In two-component granules, YTHDF1 acts as a kinetic inhibitor that attenuates YTHDF2 assembly (Fig. 4a), whereas YTHDF3 functions as a kinetic activator that promotes YTHDF2 assembly (Fig. 4b).

Within the YTHDF protein family, the conserved residue W432—which mediates m^6^A recognition—plays a critical role in regulating both the assembly kinetics of YTHDF RNP granules (Fig. 1e) and the formation of RNA-RNA interaction architectures (Supplementary Fig. 9d-e). We also find that the identity of the LCD is a key determinant of mesoscale assembly kinetics and underlies the cross-scale emergent behaviors observed within the YTHDF protein family (Fig. 1d and 3a(iii)).

The molecular grammar of RNA m^6^A modification within RNP granules, as revealed in this study, emerges from cross-scale integration of nanometer-scale biomolecular interactions into mesoscale condensate organization. This framework suggests two general principles that warrant further investigation.

First, RNA modifications encode a molecular grammar that governs RNP granule assembly. Despite the low abundance of m^6^A and the relatively modest changes it induces in RBP binding affinity, variation in the m^6^A ratio can produce substantial regulatory effects through higher-order, cooperative assembly processes. These findings indicate that subtle differences at the nanometer scale can be amplified during the transition to mesoscale organization. Notably, RNAs harbor more than 170 distinct chemical modifications, many of which have been implicated in condensate regulation. For example, under stress conditions, *N*^1^-methyladenosine (m^1^A)-modified RNAs are enriched in stress granules ^25^, whereas 5-methylcytosine (m^5^C), one of the earliest identified RNA modifications, promotes biomolecular condensate formation of its cognate reader protein Y-box binding protein 2 ^26^. We therefore propose that diverse RNA modifications may follow a common molecular grammar analogous to that defined here for m^6^A, which warrants further investigation.

Second is a central question that concerns whether YTHDF1, YTHDF2, and YTHDF3 function redundantly or exert distinct biological roles. Although their highly similar sequences and physicochemical properties have led to the proposal that they act as redundant mediators of mRNA decay ^11,27^, alternative models assign divergent functions in translation and degradation ^4^. Our SMART-epi measurements resolve this debate by demonstrating that YTHDF proteins exhibit distinct assembly kinetics under physiological m^6^A conditions, supporting functional diversification. This behavior is reminiscent of degenerate states in the hydrogen atom ^28^, where multiple quantum states share identical energies due to high symmetry but diverge upon symmetry breaking, such as Zeeman effect by applying a magnetic field. By analogy, YTHDF proteins may appear functionally equivalent at the nanometer scale, yet cross-scale assembly processes—from molecular interactions to mesoscale condensate organization—act as symmetry-breaking mechanisms that differentiate their functional outputs. We propose that this represents a form of biomolecular cross-scale degeneracy, in which apparent molecular redundancy gives rise to emergent functional diversity through higher-order assembly. Such a framework provides a conceptual basis for understanding how subtle molecular features are amplified into distinct biological functions in biomolecular condensates. Exploring this biomolecular cross-scale degeneracy represents another promising avenue for future research.

Deciphering the biological functions of complex systems requires quantitative characterization of assembly and disassembly kinetics across multiple length scales. Elucidating the principles governing RNA modification-dependent organization thus represents a central challenge and a defining frontier in molecular biology. Here, we establish SMART-epi as a generalizable platform for probing epitranscriptomic regulation. By enabling quantitative, cross-scale kinetic measurements, this approach provides a framework for linking molecular interactions to mesoscale organization and emergent function in RNP granules. More broadly, our results offer a conceptual and methodological foundation for understanding how RNA modifications encode biological function, with potential implications for therapeutic intervention.

## Author Contributions

M.X. prepared biological samples, designed and conducted biochemical and single-molecule experiments, performed data analysis. Y.L. assisted M.X. T.W. and J.L. helped to accurately calibrate the molar ratio of [m^6^A] : [A]. Z.Q. supervised the project, experimental designs, and data analysis, and wrote the manuscript with input from all authors.

## Acknowledgments

We thank the Peking Nanofab for process support. We thank the contributions of the Engineering Research Center for Semiconductor Integrated Technology, Institute of Semiconductors, Chinese Academy of Sciences. We thank Pilong Li laboratory (Tsinghua University, China) for their generous support of the plasmid of YTHDF1, YTHDF2, and YTHDF3. We thank Dr. Pilong Li and Chunlai Chen (Tsinghua University), and the members of the Z.Q. laboratory for comments on the manuscript.

